# Human airway organoids as a versatile model to study BSL-4 virus replication and pathogenesis

**DOI:** 10.1101/2025.03.20.644281

**Authors:** Joo-Hee Wälzlein, Sebastian Reusch, Jenny Ospina-Garcia, Ruth Olmer, Marc A. Schneider, Laura V. Klotz, Christian Klotz, Susann Kummer

## Abstract

Research with BSL-4 viruses such as Ebola, Marburg, and Nipah presents significant challenges due to their high virulence and the stringent containment measures required. A major limitation in studying viral pathogenesis and developing therapeutic strategies is the absence of suitable animal models that accurately replicate human disease. In this context, 3D cell culture systems offer significant advantages over traditional 2D monolayer cultures, mimicking native physiological conditions including cell polarization and composition. Human airway organoids, derived from pluripotent or adult stem cells, closely replicate the structure and function of the human respiratory system, providing a relevant and accessible environment for studying viral replication and pathogenesis. In contrast to conventional cell lines, airway organoids enable investigation of virus-host interactions within a human tissue context, providing insights that are more directly translatable to human disease. In our study, we generated airway organoids from both clinical donor tissues and commercially available nasal epithelial cells and showed in comparative analyses with whole lung tissue that these organoids are comparable in terms of cell composition. Despite donor-specific variations due to genetic factors, airway organoids derived from different sources and donors exhibit a remarkably similar cellular make-up. We further demonstrated that organoids derived from nasal swabs can effectively replicate BSL-4 viruses, establishing them as a standardized 3D model for broader research applications and advancing our understanding of these pathogens, especially in the absence of reliable animal models.

**Author Summary:** This study establishes human airway organoids as a robust model for investigating BSL-4 pathogens, such as Ebola, Marburg, and Nipah virus. Airway organoids represent reliable systems due to their ability to replicate the complexity of human respiratory epithelia and support viral infection. These organoids exhibit high susceptibility to these viruses, allowing for subsequent analysis of infection kinetics, immune evasion, and tissue-specific tropism within a controlled environment. This platform provides a powerful tool for antiviral testing and studying virus-host interactions, thus helping bridge critical gaps in high-containment virus research.

## Introduction

A significant challenge in studying BSL-4 viruses, such as Ebola, Marburg, and Nipah, is the absence of animal models that accurately replicate the disease progression observed in humans. This limitation hinders our ability to fully understand their pathogenesis and develop effective therapies for these highly virulent pathogens. Ebola virus causes severe hemorrhagic fever with high mortality rates, primarily in sub-Saharan Africa. It spreads through direct contact with body fluids and can lead to widespread outbreaks, as seen during the 2014-2016 epidemic in West Africa (Ebola Situation Report 30th March, 2016)). Marburg virus, a close relative of Ebola, also causes hemorrhagic fever with similar transmission and clinical outcomes. Both viruses belong to the Filoviridae family and are known for their rapid disease progression and high case fatality rates [1]. In contrast, Nipah virus, a member of the Paramyxoviridae family, causes encephalitis and respiratory illness. First identified in Malaysia in 1998, Nipah virus has since caused multiple outbreaks across South and Southeast Asia [2-5]. These zoonotic viruses are typically transmitted to humans from animals such as fruit bats or pigs. Currently, there are no specific treatments or vaccines available (with exception of the Ebola vaccine, which is only effective against the Ebola Zaire strain) [6-8].

Given the lack of suitable animal models, human stem cell-derived organoids offer a promising alternative for studying these viruses in a controlled environment that closely mimics human tissue [9]. This approach not only allows for the examination of virus-host interactions, but also facilitates the testing of potential therapeutic interventions, providing insights that are more predictive of human outcomes. To provide proof-of-concept, we compared several epithelial organoids derived from respiratory tissue to evaluate cell type conformity using standardized culture protocols and demonstrate the utility of these cultures for studying BSL-4 viruses in a Biosafety Level 4 environment.

Organoids are three-dimensional, self-organizing structures derived from stem cells that better replicate organ physiology compared to 2D or conventional 3D cell cultures using immortalized cell lines [10, 11]. While pluripotent stem cell-derived organoids mimic organogenesis, providing high-fidelity models of pathophysiological complexity in organ development, adult stem cell-derived organoids retain organ identity and genetic stability over time [1, 12]. Culture protocols for adult stem cell derived organoids are relatively straightforward and can easily be established in conventional cell culture laboratories. Moreover, transcriptomic studies show that adult stem cell-derived organoids exhibit gene expression profiles closely resembling their tissue of origin, particularly for airway organoids, which closely align with primary lung tissue profiles [13, 14]. This tissue-specific similarity makes them valuable models for studying organ-specific pathologies.

## Results

Airway organoids from various source material, from donor 1, 2 (both healthy lung tissue) and HNEpCs (commercial nasal epithelial cells) exhibited similar morphology and cellular marker expression typical of human respiratory epithelial tissue (**Fig 1**). Cystic and solid structures were observed, with cystic formations, mostly larger than solid structures, characterized by cavities filled with mucus or cell debris becoming larger over time (**Fig 1A-C**). Nasal epithelial cell-derived organoids predominantly formed smaller, solid structures, with some cystic formations appearing in long-term cultures (**Fig 1C**). Immunofluorescence (IFA) and qPCR analysis confirmed the presence and localization of basal cells (keratin 5, KRT5), goblet cells (mucin 5AC, MUC5AC), ciliated cells (acetylated tubulin, AcTUB), and club cells (secretoglobin family 1A member 1, SCGB1A1) in all organoids, consistent with primary airway tissue (**Fig 1D-E; Fig 2** [15]). Specifically, IFA of donor 2 and HNEpC-derived organoids revealed comparable staining for all markers. Comprehensive qPCR analysis verified the expression of all major airway epithelial cell types, with levels generally comparable to or exceeding those observed in lung tissue (**Fig 2**). Although some single marker genes showed significant donor-related variation, a notable difference between nasal and lung-derived organoids was observed in markers for ciliated cells (*FOXJ1, SNTN*). This was expected as ciliated cells are more abundant in nasal epithelial tissue in comparison to lower airway epithelia [16]. Overall, the analysis demonstrated that airway organoids derived from different sources exhibit consistent cellular characteristics and mimic human airway tissue effectively. Although nasal organoids displayed smaller lumens and a higher proportion of ciliated cells compared to lung-derived organoids, they showed stable comparable cellular composition.

**Figure 1:**
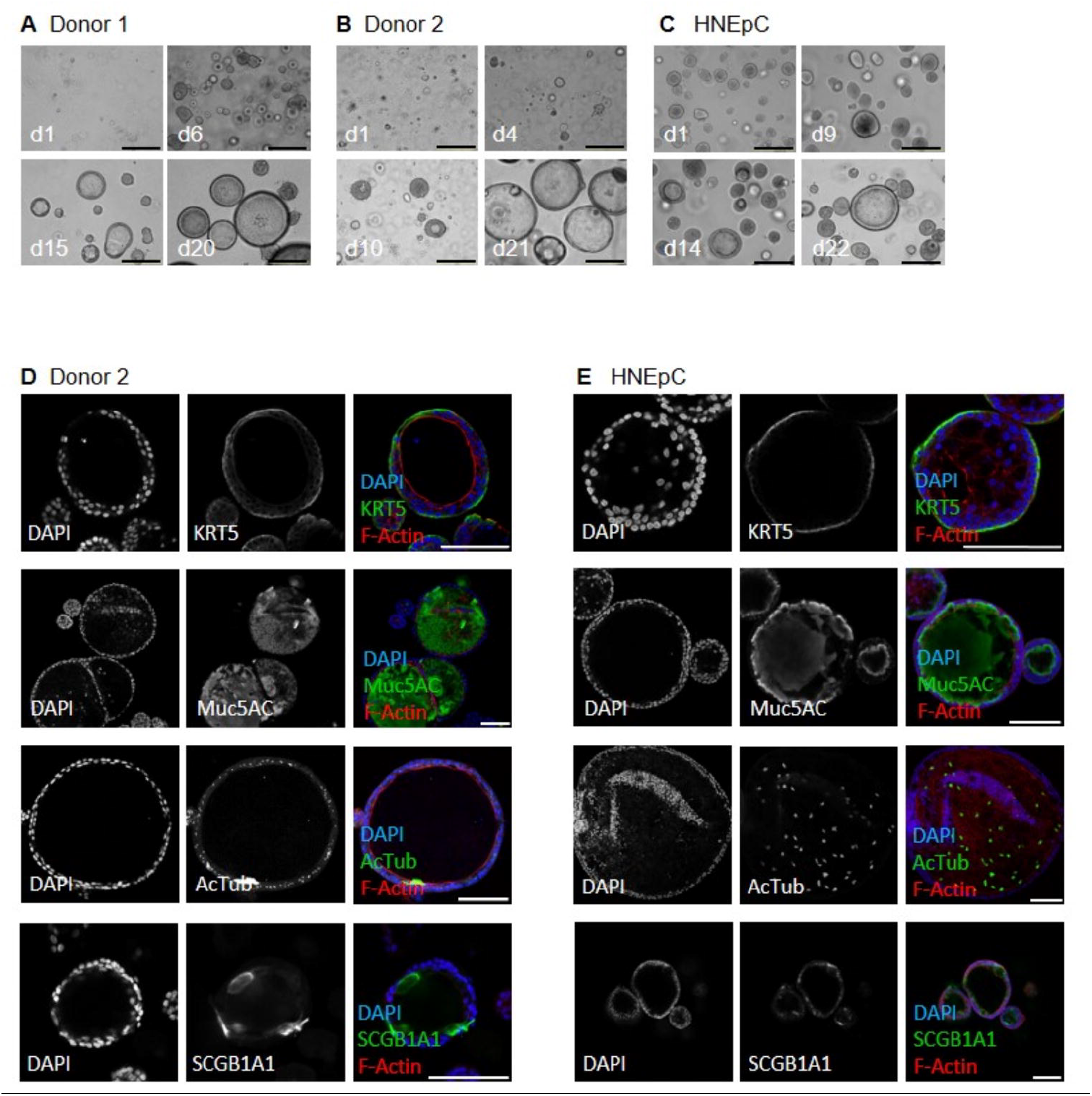
Microscopic analysis of human airway organoids (AOs) focusing on their morphology and the spatial localisation of cellular marker proteins. A-C: Bright-field images of AOs at different days of cultivation as indicated. Scale bars = 50 μm. Images were taken employing a widefield microscope and a 20x air objective (d = day). **D/E**: Immunofluorescence staining of AOs derived from adult stem cells of human lung tissue from donor 2 or HNEpC using antibodies specific to keratin5 (KRT5), mucin5AC (Muc5AC), acetylated tubulin (AcTub), and uteroglobin (SCGB1A1). Cell nuclei were counterstained with DAPI and F-actin with phalloidin, respectively. Image acquisition was performed using a confocal laser scanning microscope and a 20x immersion oil objective. Scale bars = 100 μm.

**Figure 2:**
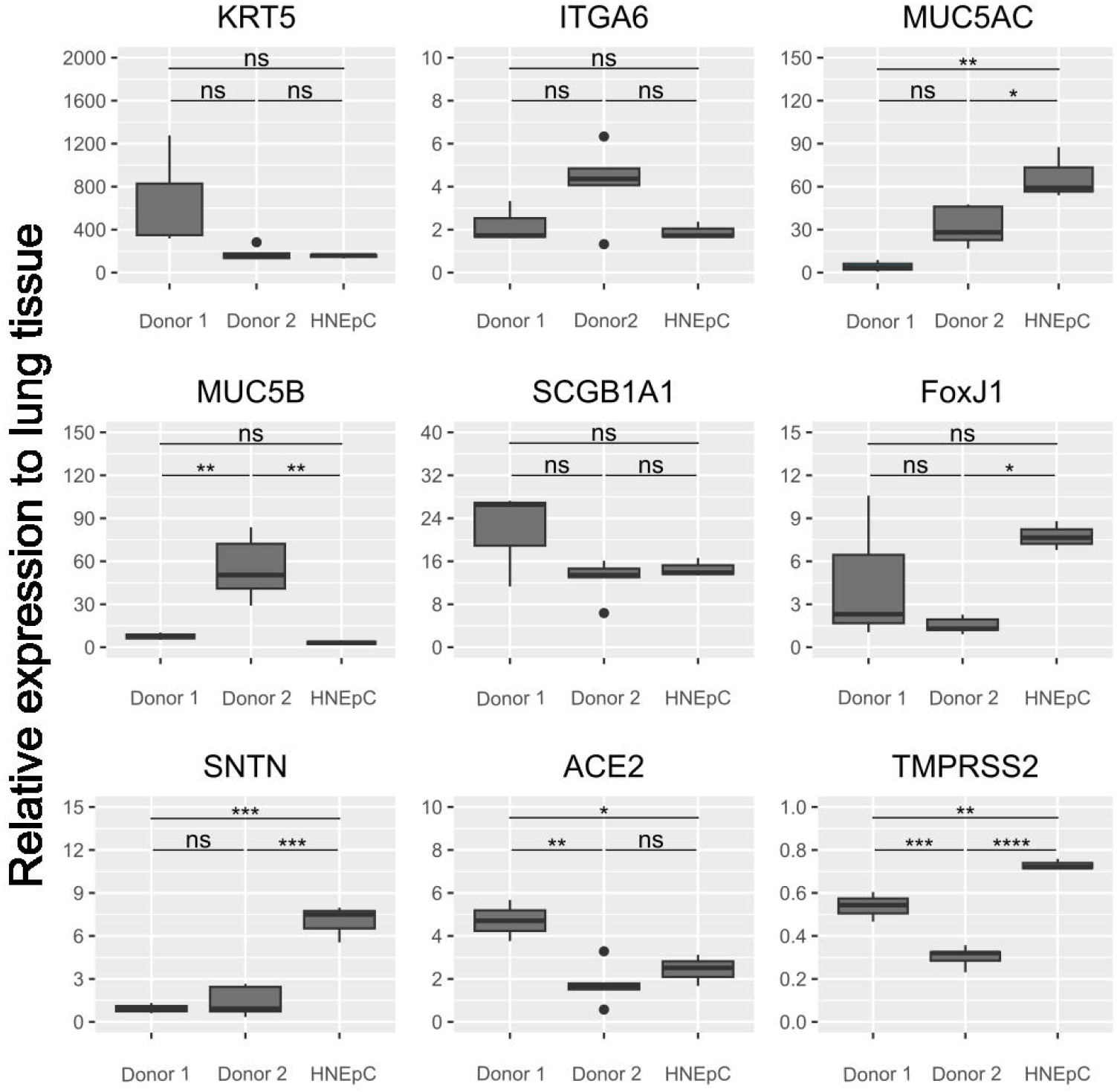
qPCR-analysis of human airway organoids (AOs) reveals expression of all major airway epithelial marker genes comparable to human lung tissue. Boxplots show the median and interquartile range, whiskers represent the highest and lowest values, outliers are plotted as dots. D1 = donor 1, D2 = donor 2, HNEpC = human nasal epithelial cells. Statistical significance was calculated using one-way ANOVA, followed by a post-hoc Tukey’s test (ns = p > 0.05; * = p ≤ 0.05; ** = p ≤ 0.01; *** = p ≤ 0.001; **** = p ≤ 0.0001).

Next, we used nasal organoids for infection studies with BSL-4 viruses. Despite their representation of smaller airways and higher abundance of ciliated cells compared to bronchial organoids, this choice was driven by their commercial availability and therefore easy accessibility. Strikingly, the nasal-derived organoids showed high susceptibility to Marburg (MARV), Nipah (NiV), and Ebola Zaire virus (EBOV). These results mark the first proof-of-concept for using these respiratory organoids to study high-risk pathogens in a controlled environment. Viral replication and release were monitored by quantifying the viral genome equivalents present within the organoids (**Fig 3**). Following incubation with the initial virus input, a notable increase in viral genome equivalents was detected, indicating effective viral replication within the nasal-derived organoids. We further investigated the presence of putative entry markers, namely Niemann-Pick disease, type C1 (NPC1) for Ebola [17] and Marburg virus and Ephrin B2 and B3 (EFNB2, EFNB3) for Nipah virus [18, 19], by qPCR analysis. Transcripts of these proteins were detected at significant levels (data not shown), leading the way to future functional studies.

**Figure 3:**
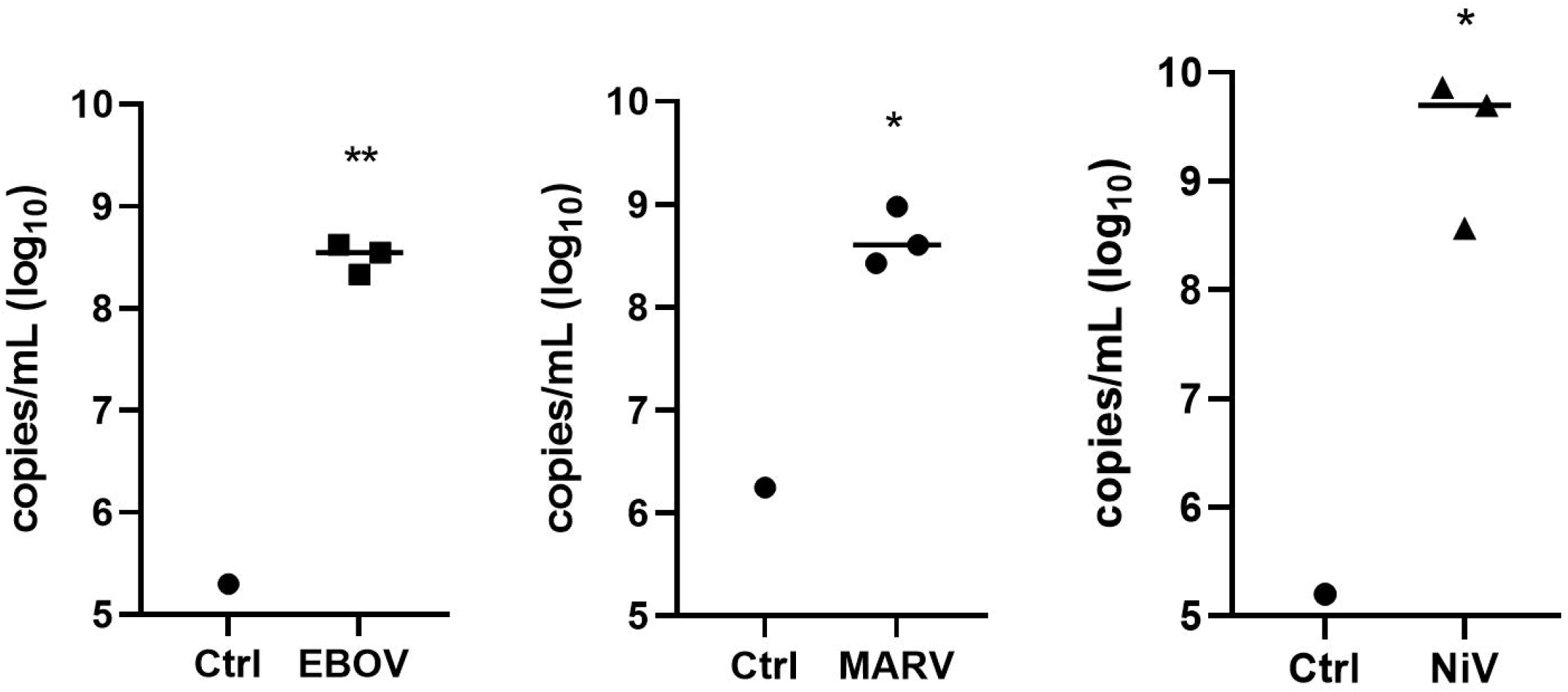
HNEpC-derived airway organoids (AOs) exhibit viral replication following infection with Ebola (EBOV), Marburg (MARV), and Nipah (NiV) virus. AOs were infected with EBOV, MARV and NiV. At 3 days post infection virus-specific copies per mL were quantified by qRT-PCR. Controls (ctrl) were exposed to the viruses for 5 min only. Statistical significance was calculated using unpaired t-test (* = p ≤ 0.05; ** = p ≤ 0.01).

## Discussion

In this study, we demonstrated that airway organoids from different origin exhibit remarkable consistency in cell type composition, yet maintain regional-specific tissue characteristics. These models are highly susceptible to infection and replication of BSL-4 viruses. By providing insights into tissue-specific tropism and viral entry mechanisms, they may help uncover the molecular determinants that enable viruses like Marburg, Nipah or Ebola virus (MARV, NiV, EBOV) to selectively infect certain tissues, such as the respiratory tract in future studies. Additionally, when supplemented with immune cells, airway organoids allow real-time analysis of immune dynamics, making them well-suited models to study immune evasion strategies of these viruses [20]. Moreover, they also represent a valuable model to study vascular endothelial interactions, helping to clarify how hemorrhagic viruses disrupt blood-tissue barriers and contribute to vascular pathology.

For therapeutic applications, airway organoids provide a robust platform to screen antiviral agents and evaluate tissue-specific drug efficacy and toxicity, with potential implications for precision medicine [21, 22]. Finally, these models facilitate exploration of virus-host interactions and virus-induced metabolic shifts, which could identify novel therapeutic targets.

The three-dimensional architecture of organoids provides an improved representation of host-virus interactions at cellular and molecular levels compared to conventional 2D cell culture systems. In the absence of suitable animal models, organoids allow for physiologically relevant analysis of BSL-4 pathogens, enhancing our understanding of complex viral behaviour and therapeutic responses. Future research should focus on refining culture protocols and enhancing physiological relevance of organoids through improved vascularization, blood perfusion, and immune cell integration.

In conclusion, our proof-of-concept study provides an easy to adapt method to study BSL-4 viruses in an advanced organoid-based airway cell culture model using commercially available cells and published, robust culture protocols.

## Material and Methods

### Patient samples

Patient-derived lung cells were provided by Lung Biobank Heidelberg, a member of the BioMaterialBank Heidelberg (BMBH) and the platform biobanking of the German Center for Lung Research (DZL). All patients signed an informed consent and study was approved by the ethics committees of Heidelberg and Berlin (S-270/2001 (biobank vote) and EA2/090/20 (study vote).

### Organoid Culture and Maintenance

Organoid cultures were established from cryopreserved lung biopsy cells following protocols from Zhou and Sachs [1,3]. Briefly, cells were thawed into DMEM/Ham F-12 medium with ROCK inhibitor, centrifuged, and seeded in organoid medium into T25 flasks. Cells were later transferred into 6-well plates with Cultrex BME for 3D culturing, and medium was replaced weekly. For passaging, cells were enzymatically dissociated with TrypLE, centrifuged, and re-embedded in Cultrex BME (see Supplement for details on culture medium and methods).

### Viruses and Infection

BSL-4 viruses, including Ebola-GFP, Ebola, Marburg, and Nipah, were obtained and titrated following standard protocols. Organoids derived from nasal epithelial cells were cultured and infected following protocols detailed in the Supplement. Infections were performed at BSL-4 containment.

### RNA Purification, cDNA Synthesis, and qPCR

Organoid RNA was extracted using the Zymo Quick-RNA Kit and converted to cDNA with the High Capacity RNA-to-cDNA Kit. qPCR was performed using both one-and two-step protocols to quantify viral RNA and analyse gene expression abundance. Primer sequences and detailed qPCR protocols are available in the Supplement.

### Immunofluorescence Staining and Microscopy

Organoids were fixed, permeabilized, and stained using primary and secondary antibodies to detect relevant markers. Image acquisition was performed on a Leica STELLARIS 8 microscope, and images were processed with ImageJ. Full antibody list and staining procedures can be found in the Supplement.

## Acknowledgments

We thank Andreas Kurth for kindly providing the possibility to perform the BSL-4 experiments using the BSL-4 containment lab at the Centre for Biological Threats and Special Pathogens, Robert Koch-Institute, Berlin, Germany. We thank Carmen Hoppstock, Nicole Kromarek, Elke Radam and Gudrun Kliem for excellent technical support.

