## Supplemental files for "Human airway organoids as a versatile model to study BSL-4 virus replication and pathogenesis"

**SUPPLEMENT**

**Material and Methods**

***Organoid cultures and maintenance***

Organoid cultures are based on established protocols outlined by Zhou and Sachs [10, 23]. Cryopreserved cells of lung biopsies containing adult stem cells (see Table 1), preserved in a 10% DMSO solution, were thawed in a 37°C water bath. Subsequently, these cells were resuspended in (a 15 mL Falcon tube containing) 5 mL of pre-warmed DMEM/Ham F-12-Gibco medium (Gibco™ CAT#12634010), supplemented with 1:1,000 Y-27632 (ROCK inhibitor; AbMole Bioscience CAT#M1817). Following centrifugation at 500 x RCF for 5 min at 4°C, the pellet was resuspended in 10 mL of organoid medium (see Table 2) and seeded in a T25 flask. Cells were incubated overnight at 37°C with 5% $CO_2$. For subsequent steps of the 3D-culture, all components were kept on ice. The following day, cells from the T25 flask were collected, adherent cells were gently detached using a cell scraper, and transferred to a 15 mL Falcon tube coated with 0.1% BSA, kept on ice. Cells were then centrifuged at 500 x RCF for 5 min at 4°C and the pellet was resuspended in an ice-cold mixture of 70% Cultrex Basement Membrane Extracts (BME, growth factor-reduced Type 2 basement membrane extract; R&D Systems CAT#3533-005-02) and 30% organoid medium. The cells were seeded as 70 µL BME droplet domes in 6-well plates, with each well containing three domes, and a total of one to three wells used, depending on the initial cell number. The plates were incubated for 30 min at 37°C to allow solidification of the BME. Once solidified, 2mL organoid medium was added to the domes and medium was changed twice a week. For subsequent passages and culture expansion, BME droplets were mechanically disrupted by washing over them with organoid medium using a P1,000 tip and then centrifuged at 500 x RCF for 5 min at 4°C. This was followed by enzymatic digestion using TrypLE Express (Gibco™ CAT#12605010), supplemented with 10 µM ROCK inhibitor for 4-8 min at 37°C. Cells were dispersed into a single-cell solution using a blunt needle (18G), followed by the addition of DMEM/Ham F-12 medium supplemented with ROCK inhibitor. After centrifugation as described earlier, cells were embedded again in 70% BME split in a 1:3 or 1:6 ratio, again in the form of droplets (each well containing three droplets). The organoids used for both characterization and infection experiments underwent two passages after thawing the cells. For each biological sample, a complete 6-well plate was utilized, comprising a total of 18 droplets.

**Table 1: Sources of adult stem cells used for airway organoid generation**

| Material | Source | Description |
|---|---|---|
| Donor 1 | Lung Biobank Heidelberg, member of the accredited Tissue Bank of the National Center for Tumor Diseases (NCT) Heidelberg, the Biomaterial Bank Heidelberg, and the Biobank platform of the German Center for Lung Research (DZL) | Lung biopsy of non-cancerous tissue from a 70-year-old, non-smoking patient diagnosed with pulmonary carcinoid |
| Donor 2 | Leibniz Research Laboratories for Biotechnology and Artificial Organs (LEBAO), Biomedical Research in Endstage and Obstructive Lung Disease Hannover (BREATH), German Center for Lung Research (DZL), Department of Cardiothoracic, Transplantation and Vascular Surgery Hannover Medical School, Hannover, Germany) | Lung biopsy of non-cancerous tissue from a 50-year-old patient diagnosed with fibrosis |
| HNEpC | PromoCell (CAT #C-12620, Lot 475Z023) | Primary human nasal epithelial cells isolated from normal human nasal mucosa |
| Lung tissue | DZL Heidelberg | |

**Table 2: Complete organoid medium**

| Medium supplement | Final concentration |
|---|---|
| Advanced DMEM/F12 | 1x |
| R-Spondin 1 | 500 ng/mL |
| FGF-7 | 25ng/mL |
| FGF-10 | 100ng/mL |
| Noggin | 100ng/mL |
| A83-01 | 500nmol/L |
| Y-27632 | 5µmol/L |
| SB202190 | 500nmol/L |
| N-Acetylcysteine | 1.25mmol/L |
| Nicotinamide | 5mmol/L |
| B27 | 1x |
| Pen/Strep/Glutamine | 100U/mL; 100µg/mL; 1x |
| Hepes | 10mmol/L |

| | |
|---|---|
| Primocin | 50µg/mL |

**Viruses and infection**

Ebola-GFP [24], Ebola virus (Makona), and Marburg virus (Musoke) were kindly provided by Prof. Dr. Stephan Becker, Institute for Virology, Philipps-University Marburg, Germany, and Nipah virus (Malaysia) was kindly provided by Heinz Feldmann, Rocky Mountain Laboratory, Hamilton, Montana, USA. All viruses were propagated in green monkey kidney epithelial Vero E6 cells (American Type Culture Collection, Manassas, VA, CAT#CCL-1586). The viral titer was determined using the Spearman and Kärber method, specifically the $TCID_{50}$/mL (Tissue Culture Infectious Dose 50 per milliliter) assay [25].

The experiments involving risk group 4 viruses were conducted at the BSL-4 facility of the Robert Koch-Institute in Berlin, Germany, strictly following standard operating protocols for handling infectious agents. Organoids derived from commercially available airway epithelial cells sourced from nasal swabs were cultured for approximately 20 d prior to infection as previously described. Organoids grown in 18 x 70 µL domes were collected into 15 mL tubes and centrifuged at 500 × RCF for 5 min, leaving a liquid pellet containing the organoids. The supernatant was discarded and BME was removed by incubating the pellet in 3 mL cell recovery solution for 45 min at 4°C with resuspension every 15 min, followed by another centrifugation step. Subsequently, 1260 µL of 70% BME, diluted in organoid medium, was mixed with 25 µl EBOV ($TCID_{50}$ = 2.94x $10^7$/mL); 18.4 µl MARV ($TCID_{50}$ = 3.98x $10^7$/mL) and 11.7 µl NiV ($TCID_{50}$ = 6.3x$10^7$/mL), respectively, to reach equal levels of infection doses; organoids were resuspended in this mix avoiding bubble formation on ice. This mixture was then seeded as droplets in 6-well LabTeks plates and incubated at 37°C for 30 min. Once solidified, the BME droplets were maintained for the specified durations in a previously reported medium [10]. Controls were treated simultaneously, but infection was stopped after 1 h. Organoids were collected and pelleted as described above in this section and organoids were re-seeded in 70% BME in the absence of viral input.

### *RNA purification, reverse transcription and qPCR*

The primer pairs (see table 3) for the different markers were validated and efficiency of primers was calculated as described [26].

**Table 3: Primer used for qRT-PCR**

| Target | Sequence | Gene ID |
|---|---|---|
| **KRT5** | Fw: CCAAGGTTGATGCACTGATGG<br><br>Rev: TGTCAGAGACATGCGTCTGC | 3852 |
| **MUC5AC** | Fw: CAGCACAACCCCTGTTTCAAA<br><br>Rev: GCGCACAGAGGATGACAGT | 4586 |
| **SCGB1A1** | Fw: TCCTCCACCATGAAACTCGC<br><br>Rev: AGGAGGGTTTCGATGACACG | 7356 |
| **FOXJ1*** | Fw: AGATCCCACCTGGCAGAATTCAA<br><br>Rev: CCGAGGCACTTTGATGAAGC | 2302 |
| **SNTN** | Fw: TGTATGCACAGTACCCAGGAC<br><br>Rev: AGCAGTGGTGGCAATAGCTTT | 132203 |
| **MUC5B** | Fw: GCCTACGAGGACTTCAACGTC<br><br>Rev: CCTTGATGACAACACGGGTGA | 727897 |
| **ITGA6*** | Fw: ATGCACGCGGATCGAGTTT<br><br>Rev: TTCCTGCTTCGTATTAACATGCT | 3655 |
| **TMPRSS2*** | Fw: CAAGTGCTCCAACTCTGGGAT<br><br>Rev: AACACACCGATTCTCGTCCTC | 7113 |
| **ACE2*** | Fw: CAAGAGCAAACGGTTGAACAC | 59272 |

| Target | Sequence | Gene ID |
|---|---|---|
| | Rev: CCAGAGCCTCTCATTGTAGTCT | |
| **GAPDH*** | Fw: CTCCTGTTCGACAGTCAGCC<br><br>Rev: CCCAATACGACCAAATCCGTTG | 2597 |
| **ß-Actin*** | Fw: CATGTACGTTGCTATCCAGGC<br><br>Rev: CTCCTTAATGTCACGCACGAT | 60 |
| **RPS18*** | Fw: AGTTCCAGCACATTTTGCGAG<br><br>Rev: TCATCCTCCGTGAGTTCTCCA | 6222 |
| **GFP** | Fw: GAGCGCACCATCTTCTTCAA<br><br>Rev: CTGCTTGTCGGCCATGATATAG | 25339618 |
| **Ebola Zaire, VP30** | Fw: ACT CCT ACT AAT CGC CCG TAA G<br>Rev: ATC AGC CGT TGG ATT TGC T<br>Probe: CACCCAA+GGACTCGC | |
| **Marburg, NP** | Fw: GTCCTCAGCCAGAAACGAGA<br>Rev: ACCGTTACTTCCACAGGTGT<br>Probe: TCACAGAATCGGGTGTCACAGTCGT | |
| **Nipah Malaysia, NP** | Fw: GTTCAGGCTAGAGAGGCAAAATTT<br>Rev: CCCCTTCATCGATATCTTGATCA<br>Probe: CTGCAGGAGGTGTGCTCATTGGAGG | |

* Sequence taken from PrimerBank, Harvard Medical School.

This study employed two distinct quantitative polymerase chain reaction (qPCR) protocols. The initial protocol, which utilized a two-step qPCR method, was applied for organoid characterization, while a one-step qPCR approach was implemented for the infection experiments. For the two-step qPCR, 18 organoid domes embedded in BME were collected, centrifuged at 500 x RCF for 5 min at 4°C. Subsequently, the pellet was resuspended in 700 µl of Trizol Reagent (Zymo Research CAT#R2050-1-200) and stored at -80°C until further processing. RNA extraction was performed using the Zymo Quick-RNA Microprep Kit (Zymo Research CAT#R1050) following the manufacturer`s instructions. RNA concentration was assessed using Tecan's NanoQuant Plate. Synthesis of cDNA was employed using the High Capacity RNA-to-cDNA kit (Applied Biosystems-Thermo Fisher Scientific CAT#4387406). 20 µL of the reaction mixture were pipetted into ice-cold RNase-free PCR tubes. During the incubation step, the mixture was heated to 70°C for 5 min without the addition of RT Enzyme Mix, facilitating denaturation and allowing for initial annealing. The RT program encompassed 60 min at 37°C, 5 min at 95°C, and a final step at 4°C. After cDNA synthesis, samples were diluted with nuclease-free water to a concentration of 2 ng/µL in preparation for qPCR analysis. Real-time quantification was carried out using 10 ng (5 µL) of cDNA with SYBR Green LUNA Universal qPCR (New England BioLabs CAT#M3003) on a Bio-Rad CFX96 qPCR real-time thermocycler. For the one-step qPCR, 18 embedded organoids were collected, centrifuged at 500 x RCF for 5 min at 4°C. BME was removed by incubating the pellet in 3 mL cell recovery solution for 45 min at 4°C with resuspension every 15 min. Following another centrifugation step the supernatant was removed and organoids were lysed in 300 µL RLT buffer and RNA was isolated following the manufacturer`s instruction (Qiagen CAT# 74104). RNA samples for one-step qPCR analysis were diluted with nuclease-free water to 2 ng/µL. Real-time quantification was performed using the SYBR Green One-Step RT-qPCR Kit (New England BioLabs CAT#E3005) with 10 ng of RNA on a Bio-Rad CFX96 qPCR Real-Time thermocycler.

***Immunofluorescence staining and laser scanning microscopy.***

Organoids were collected and centrifuged at 500 x RCF for 5 min at 4°C. Thereafter, in order to prevent organoid damage, centrifugation was avoided and, cells were allowed to settle naturally. The pellet containing organoids and BME were incubated in cell recovery solution (Corning CAT#CLS354253) for 45 min at 4°C to dissolve the extracellular matrix. During the incubation organoids were diligently resuspended every 10 min using a 0.1% BSA-coated pipette tip. After settling, supernatant was aspirated and the pellet was washed with PBS to eliminate any residual supernatant. Subsequently, the organoids were fixed in 4% paraformaldehyde (PFA, ROTH CAT#0335.1) for 45 min at 4°C. During the fixation process, organoids were gently resuspended every 10 min with a BSA-coated pipette tip. Following fixation, two additional PBS washes were performed. The organoids were then incubated in permeabilization buffer (1xTBS, 0.25% TritonX-100, 0.1 M glycine for 20 min) at room temperature.

Following settling, the permeabilization buffer was carefully removed and samples were incubated in blocking buffer (1xTBST, 0.02% Triton-X100, 3% BSA, 1% normal goat serum) for 1 h at room temperature. Primary antibodies (see Table 4) were diluted in 900 µl of blocking buffer as indicated. The organoids were incubated overnight with the primary antibodies at 4°C, washed three times with washing buffer (1xTBS-T containing 0.05% Tween and 0.02% Triton-X100) and allowed to settle before removing the supernatant. Afterwards, the organoids were incubated with the secondary antibodies (see Table 4) in blocking buffer, DAPI for nuclear staining, and phalloidin for actin filament staining at room temperature for approximately 2 h. Following staining, three washes with washing buffer, as described above, were carried out, followed by a final wash with Milli-Q water. Finally, the organoids were gently resuspended in approximately six to eight drops of non-hardening mounting medium (Ibidi, CAT#50001) and placed onto microscope slides. During the mounting process, spaces were left between the slide and the coverslip to prevent organoid compression. These spaces were sealed with nail polish to ensure proper preservation.

Image acquisition was performed using an inverted fluorescence confocal microscope (Leica STELLARIS 8) with a 20x objective. The imaging software employed was LAS X, with image resolution set to 1024 × 1024 pixels, and subsequent image editing was conducted using ImageJ software.

**Table 4: Primary and secondary antibodies employed for immunofluorescence staining**

| **Antibody** | **Company** | **Order Number** | **Dilution** | **Host** |
|---|---|---|---|---|
| ***Primary Antibodies*** | | | | |
| Anti-CC10-E11 | Santa Cruz | sc365992 | 10ug/ul | Mouse |
| Anti-Mucin 5AC | Thermo Fisher | MA5-12178 | 1:100 | Mouse |
| Anti-acetylated tubulin | Sigma Aldrich | T6793 | 1:100 | Mouse |
| Anti-Cytokeratin 5 | Biolegend | 905501 | 1:100 | Rabbit |
| ***Secondary Antibodies*** | | | | |

| | | | | |
|---|---|---|---|---|
| αMouse IgG (H+L) | Thermo Fisher (Alexa Fluor647) | A28181 | 1:250 | Goat |
| αRabbit IgG (H+L) | Thermo Fisher (Alexa Fluor647) | A27040 | 1:500 | Goat |
| Phalloidin iFluor | Abcam (iFluor 488) | ab176753 | 1:1,000 | - |
| DAPI | Abcam | ab228549 | 1:500 | - |

***Data analysis***

The qRT-PCR data were subjected to analysis using the ΔΔCt method [6]. For assessing the expression changes of the gene of interest (GOI), the mean Cq values of the biological replicates for the GOI were initially computed. These Cq values were then normalised against the reference gene ß-Actin. The obtained results from the control (lung tissue) and donor samples were juxtaposed, and ΔΔCq values were computed.

Statistical analysis was performed using Prism9 software (GraphPad). Ordinary one-way ANOVA was employed for qPCR analyses pertaining to organoid characterization, comparing means across more than two populations, assuming normal distribution and equal variance for all tests. Boxplots show the median and interquartile range, whiskers represent the highest and lowest values, outliers are plotted as dots. For analysis of qPCR data showing viral replication, unpaired t-test was performed. The presented p-values are as follows: ns (not significant) = $p > 0.05$; * = $p \leq 0.05$; ** = $p \leq 0.01$; *** = $p \leq 0.001$; **** = $p \leq 0.0001$.
